# Cell aggregation and aerobic respiration facilitate survival of *Zymomonas mobilis* ZM4 in an aerobic minimal medium

**DOI:** 10.1101/526699

**Authors:** Sara E. Jones-Burrage, Timothy A. Kremer, James B. McKinlay

## Abstract

*Zymomonas mobilis* produces ethanol from glucose near the theoretical maximum yield, making it a potential alternative to yeast for industrial ethanol production. A potentially useful industrial feature is the ability to form multicellular aggregates called flocs, which can settle quickly and exhibit higher resistance to harmful chemicals. While spontaneous floc-forming *Z. mobilis* mutants have been described, little is known about the natural conditions that induce *Z. mobilis* floc formation and the genetic factors involved. Here we found that wild-type *Z. mobilis* forms flocs in response to aerobic growth conditions but only in a minimal medium. We identified a cellulose synthase gene cluster and a single diguanylate cyclase that are essential for both floc formation and survival in an aerobic minimal medium. We also found that NADH dehydrogenase 2, a key component of the aerobic respiratory chain, is important for survival in an aerobic minimal medium, providing a physiological role for this enzyme which has previously been found to be disadvantageous in aerobic rich media. Supplementation of the minimal medium with vitamins also promoted survival but did not inhibit floc formation.

**Importance:** The bacterium *Zymomonas mobilis* is best known for its anaerobic fermentative lifestyle, in which it converts glucose into ethanol at a yield that surpasses that of yeast. However, *Z. mobilis* also has an aerobic lifestyle which has confounded researchers with attributes of poor growth, accumulation of toxic acetic acid and acetaldehyde, and respiratory enzymes that are detrimental for aerobic growth. Here we show that a major *Z. mobilis* respiratory enzyme and the ability to form multicellular aggregates, called flocs, are important for survival but only during aerobic growth in a medium containing a minimum set of nutrients required for growth. Supplements like vitamins or yeast extract promote aerobic growth, and in some cases inhibit floc formation. We propose that *Z. mobilis* likely requires aerobic respiration and floc formation to survive in natural environments that lack protective factors found in supplements like yeast extract.

## Introduction

*Zymomonas mobilis* is a bacterium that can naturally produce ethanol from glucose near the theoretical maximum yield (1). Due to this high ethanol yield, *Z. mobilis* is often viewed as the bacterial counterpart to the yeast *Saccharomoyces cerevisiae* (2, 3), although *Z. mobilis* has yet to be widely adopted for ethanol production on an industrial scale. Several physiological differences distinguish these two ethanol producers. For example, to metabolize sugar, *Z. mobilis* uses the low-ATP yielding Entner-Doudoroff pathway whereas *S. cerevisiae* uses the more energetically efficient Embden-Myerhof-Parnas pathway. *Z. mobilis* can also use inexpensive N_2_ gas as the sole nitrogen source without compromising the high ethanol yield (4) whereas yeast, like all eukaryotes, cannot use N_2_. These two ethanol producers also differ in their ability to tolerate inhibitory compounds in feedstock hydrolysates. For example, *S. cerevisiae* shows a greater tolerance of soy hydrolysate (5) and *Z. mobilis* shows a greater tolerance of drought-stressed switchgrass hydrolysate (6).

Another difference between these two ethanol producers is the tendency of *S. cerevisiae* to rapidly settle out of solution, in some cases aided by the formation of multicellular clusters called flocs (7). Flocculation can be advantageous for collecting cells or for immobilizing cells for use in continuous flow bioreactors (7). The genetic factors behind *S. cerevisiae* flocculation are relatively well understood (8, 9) to the extent that flocculation can be genetically manipulated (10). Flocs have also been observed in *Z. mobilis* and have been studied after enriching for floc-forming mutants that arouse spontaneously (11) or after chemical mutagenesis (12). Research on *Z. mobilis* flocs has thus far primarily focused on their utility in cell-recycle bioreactors (12) and their increased tolerance of inhibitory chemicals found in cellulosic hydrolysates (13). However, relatively little physiological and genetic characterization of *Z. mobilis* flocs has been performed beyond determining that cells within flocs are held together by an extracellular matrix containing cellulose (11, 14, 15). A gene cluster likely involved in cellulose synthesis was also recently identified (15).

Here we describe physiological conditions and genetic factors that are required for floc formation in wild-type *Z. mobilis* ZM4. We found that flocs form in response to aerobic conditions but only when cultured in a minimal medium. We verified the involvement of a cellulose synthase gene cluster and additionally identified a single diguanylate cyclase (DGC) that is required for flocculation. We found that flocculation is required for cell viability in an aerobic minimal medium, suggesting a protective role against some aspect of aerobic metabolism. Furthermore, we found that NADH dehydrogenase 2 (NADH DH), a key enzyme in aerobic respiration, is important for survival in an aerobic minimal medium, providing a physiological role for this enzyme which has otherwise been found to be detrimental for growth in aerobic rich media (16–18).

### Results

#### *Z. mobilis* ZM4 forms flocs in an aerobic minimal medium

Although best known for fermentative production of ethanol under anaerobic conditions, *Z. mobilis* is also capable of aerobic respiration (19). When we grew wild-type *Z. mobilis* ZM4, hereon referred to as ZM4, under aerobic conditions in *Zymomonas* minimal medium with NH_4_Cl as the sole nitrogen source (ZYMM), we observed that ZM4 formed flocs, visible to the naked eye (Fig. 1A). Electron microscopy verified that the flocs were multicellular aggregates (Fig. 1B). Flocs were not observed when ZM4 was grown aerobically in the undefined peptone-yeast extract-glucose (PYG) medium (Fig. 1C). Thus, aerobic conditions alone are insufficient to stimulate floc formation. Flocs were also not observed when ZM4 was grown anaerobically in either ZYMM (Fig. 1D) or in PYG (Fig. 1E). Thus, flocs do not form in response to some component of ZYMM. Rather floc formation appears to be stimulated by a combination of aerobic conditions and growth in a minimal medium.

**Fig. 1.**
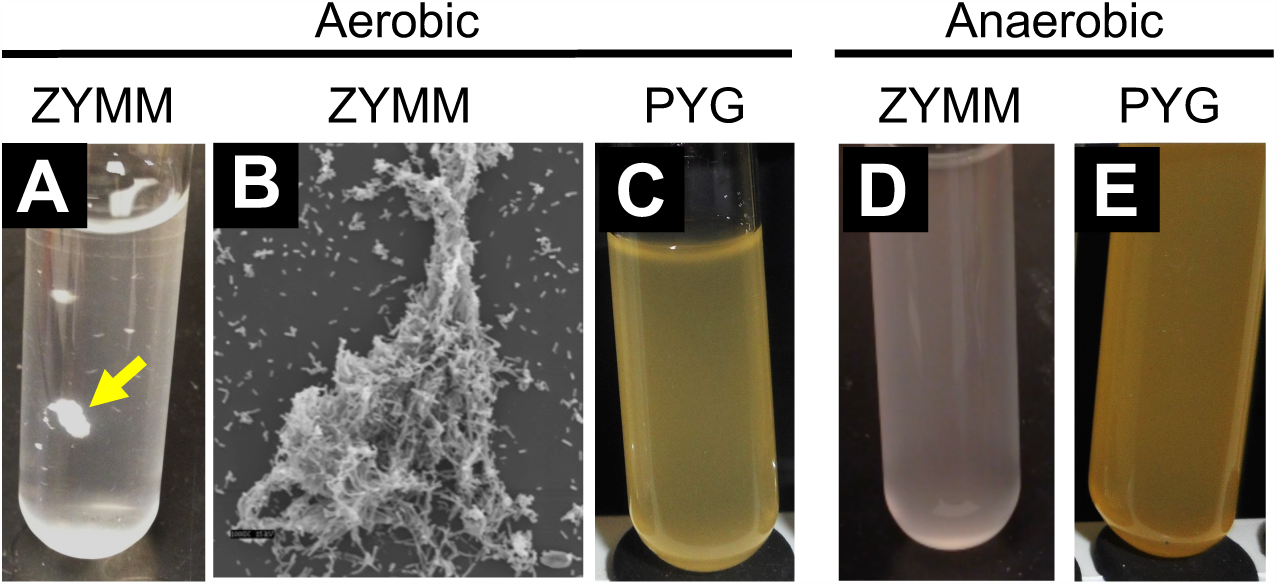
Flocs are only observed in an aerobic minimal medium. (A, C-E) Photos of *Z. mobilis* ZM4 cultures in test tubes with either a minimal medium, (ZYMM), or in undefined rich PYG medium. Media were swirled by hand prior to taking a photo. (A) The yellow arrow points to a macroscopic floc. (B) Scanning electron micrograph of a floc grown in aerobic ZYMM with NH_4_Cl. Magnification is 1000X at 15 kV.

#### Flocculation can be quantified through an increase in turbidity upon disruption

Flocs settle to the bottom of test tubes but are easily observed after swirling the tube (Fig. 1A). Flocs then settle again within seconds. To quantify the extent of flocculation, we reasoned that disruption of flocs should result in an increase in culture turbidity. Previous work by others on flocculant isolates of *Z. mobilis* determined that cellulose is a major component of the extracellular matrix; addition of cellulase to floc-forming isolates dispersed flocs whereas amylases, dextranase, pectinase, and hemicellulase did not (11, 14, 15). We verified that ZM4 flocs can be disrupted by cellulase resulting in an increase in culture turbidity (Fig. 2). Protein and DNA can also be components of an extracellular matrix for some bacteria. However, we found that neither DNAse nor Proteinase K disrupted ZM4 flocs, nor did these enzymes enhance disruption of flocs when combined with cellulase (Fig. 2A). Adding cellulase and proteinase K together decreased the rate of turbidity increase, likely due to proteinase degrading cellulase (Fig. 2A). Our findings support those of another group, which similarly found that protease did not disrupt *Z. mobilis* flocs (15). Thus, the addition of cellulase alone can be used to quantify ZM4 flocculation by an increase in culture turbidity.

**Fig. 2.**
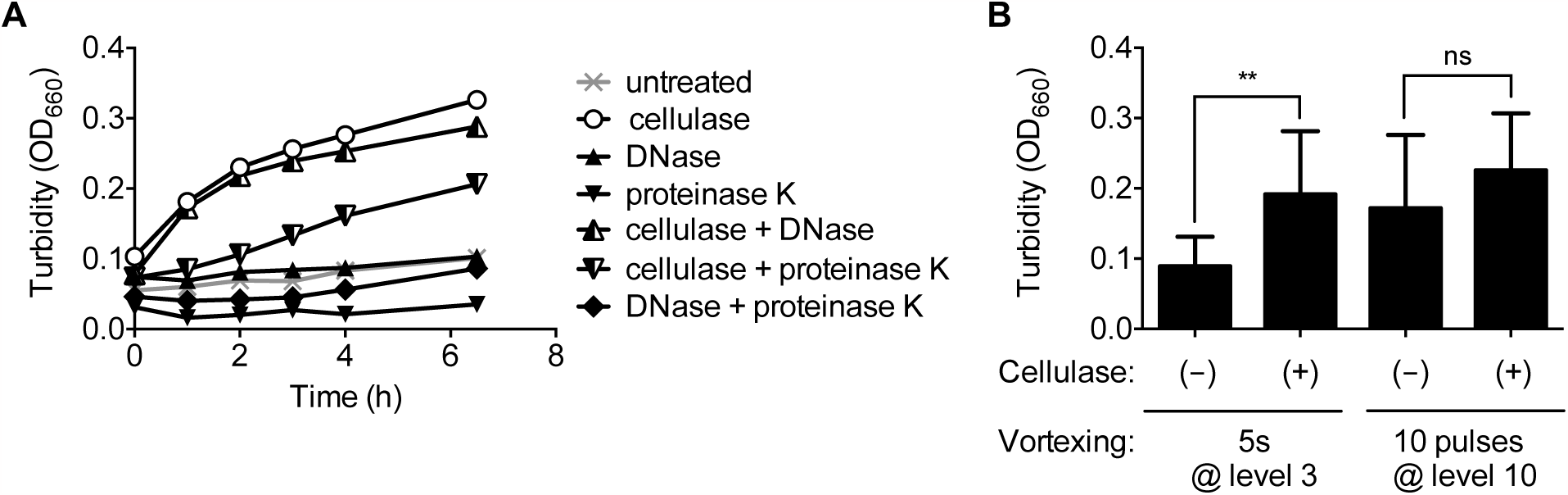
Flocs can be disrupted by cellulase or harsh vortexing. (A) Treatment of ZM4 flocs with cellulase, proteinase K, and DNase. Representative trends from one of two replicates is shown.(B) Effect of gentle (5 seconds at level 3) or harsh vortexing (10 pulses at level 10) on floc dispersal with and without a 3 h cellulase treatment. Error bars, SD (n = 4). ** Significant difference (P-value < 0.01) from a one-way ANOVA with Holm-Sidak’s posttest; ns, no significant difference.

We also found that flocs could be mechanically disrupted by harsh vortexing. Vortexing at level 3 for 5 seconds, which we refer to as gentle vortexing, briefly lifted flocs off the bottom of a test tube before they rapidly settled again. Vortexing with 10 pulses at maximum speed, which we refer to as harsh vortexing, dispersed flocs and resulted in an increase in culture turbidity that was statistically similar to that observed by adding cellulase (Fig. 2B). We used these two approaches to assess floc formation moving forward in this study.

#### A cellulose synthase gene cluster and a single diguanylate cyclase are required for flocculation

Because the extracellular matrix of flocs contains cellulose we examined the ZM4 genome sequence for genes that could be responsible for cellulose synthesis. A gene cluster from ZMO1083-1086 was annotated to encode a cellulose synthase (Fig. 3A). ZMO1086 had previously been purified and its endoglucanase activity characterized (20). More recently, another group found that genetic disruption of ZMO1083 prevented floc formation in a constitutively flocculant *Z. mobilis* mutant (15). We tested the ability of three ZM4 mutants, each containing a transposon (Tn) insertion in a different gene in this cluster, for their ability to form flocs in aerobic ZYMM. None of the mutants formed flocs in this aerobic minimal medium (Fig. 3B). We chose to complement the 1086::Tn mutant since it is the terminal gene in the operon and thus is the least likely to have polar effects from the Tn insertion. Expressing ZMO1086 from its native promoter on a plasmid restored floc formation to the 1086::Tn mutant, whereas the mutant with an empty vector failed to form flocs (Fig. 3B). We concluded that this gene cluster is required for floc formation, most likely through the production of extracellular cellulose.

**Fig. 3.**
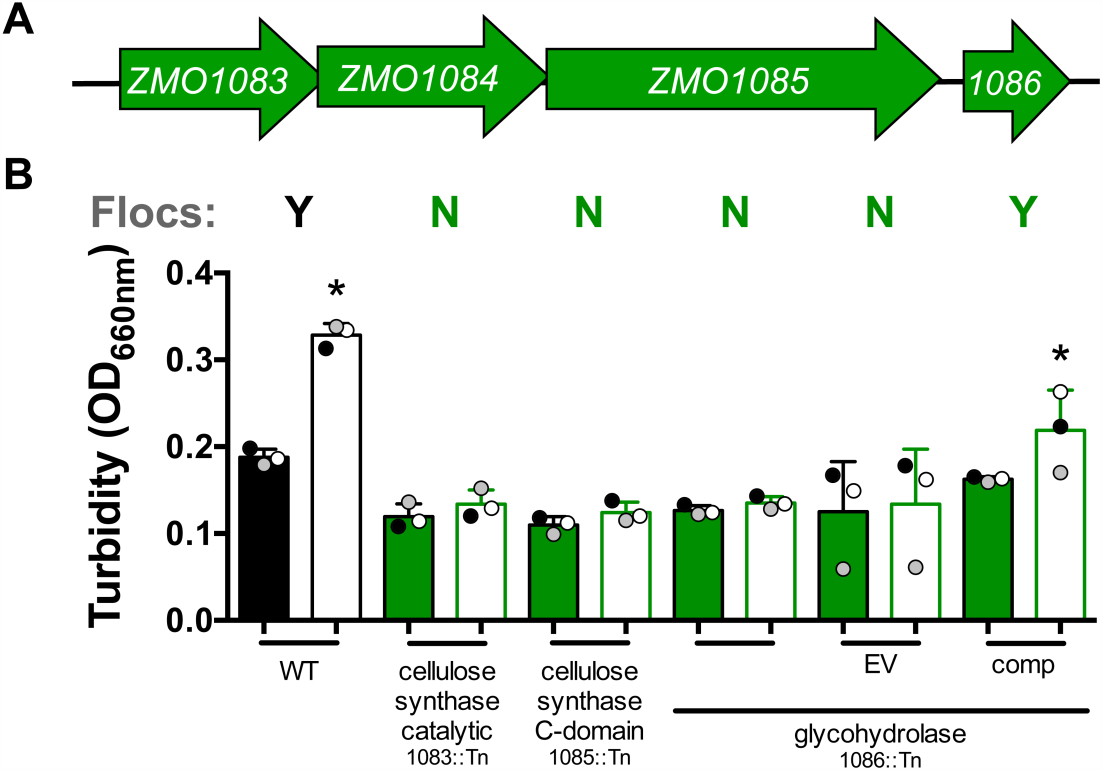
Floc formation requires cellulose synthase. (A) Cellulose synthase gene cluster (ZMO1083-1086). (B) Turbidity of aerobic cultures grown from colonies in a minimal medium (ZYMM) before (filled) and after (open) cellulase treatment. Shaded circles indicate the same culture pre- and post-disruption. Y/N indicates visual observation of flocs (Y) or not (N). WT, wild-type ZM4; EV, empty vector pSRKT; comp, complementation vector pSRKTc-1086. Error bars, SD. *Significant difference from the corresponding filled bar (P-value < 0.1) from a one-way ANOVA with Sidak’s multiple comparison posttest.

Clustering of cells into flocs or biofilms is commonly coordinated by the intracellular levels of cyclic-diguanylate monophosphate (c-diGMP) (21). c-diGMP is produced by DGCs, typically having a GG(D/E)EF domain and is degrade by phosphodiesterases (PDE), often having an EAL domain (22). The ZM4 genome has five genes annotated to have GG(D/E)EF and/or EAL domains (Fig. 4A). We examined a Tn mutant for each of the five genes for flocculation in ZYMM. Only a Tn insertion in ZMO0919, a predicted DGC, prevented floc formation (Fig. 4B). Floc formation was restored in the 0919::Tn mutant by expressing ZMO0919 under its native promoter from a plasmid, whereas an empty vector did not restore flocculation (Fig. 4B). Based on these observations, we conclude that ZMO0919 is required for floc formation, likely through the production of c-diGMP.

**Fig. 4.**
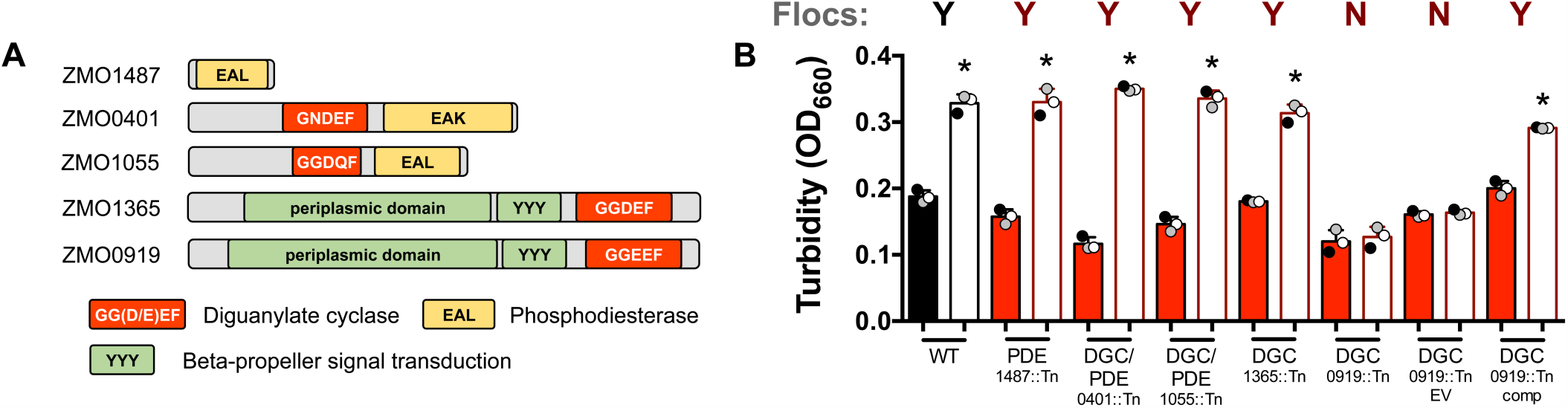
Floc formation requires cellulose synthase and a specific diguanylate cyclase. (A) Domains of ZM4 proteins with predicted diguanylate cyclase GG(D/E)EF domains (red) or phosphodiesterase EAL domains (yellow). (B) Turbidity of aerobic cultures grown from colonies in a minimal medium (ZYMM) before (filled) and after (open) cellulase treatment. Shaded circles indicate the same culture pre- and post-disruption. Y/N indicates visual observation of flocs (Y) or not (N). WT, wild-type ZM4; EV, empty vector pSRKT; comp, complementation vector pSRKTc-0919. Error bars, SD. *Significant difference from the corresponding filled bar (P-value < 0.1) from a one-way ANOVA with Sidak’s multiple comparison posttest.

#### Mutants incapable of floc formation have low viability in an aerobic defined medium

Mutants that did not form flocs in the aerobic minimal medium also had final optical densities that were lower than those of disrupted ZM4 cultures (Fig. 3B and 4B). This discrepancy in final OD values was exaggerated when mutant cultures were transferred to fresh ZYMM (Fig. 5A). These observations suggested that flocs are important for survival in aerobic ZYMM. Indeed, 1086::Tn mutant viable cell counts were below the detection limit (~1000 CFU/ml) after being transferred and incubated in fresh ZYMM, unless ZMO1086 was expressed from a plasmid (Fig. 5B). In a separate experiment, we also plated the 1086::Tn mutant for CFUs immediately after vortexing flocs in cultures that had grown from colonies in aerobic ZYMM. CFUs only developed for ZM4 and the 1086::Tn complemented strain but not for the 1086::Tn mutant with and without empty vector (data not shown; detection limit of 100 CFU/ml). Thus, the 1086 mutant was not viable even before transfer to fresh aerobic medium. Turbid growth of the 1086::Tn mutant was indistinguishable from that of ZM4 when grown from colonies in anaerobic ZYMM or in aerobic PYG (Fig. 5C). Thus, the poor growth of the 1086::Tn mutant in aerobic ZYMM was not due to a general growth defect.

**Fig. 5.**
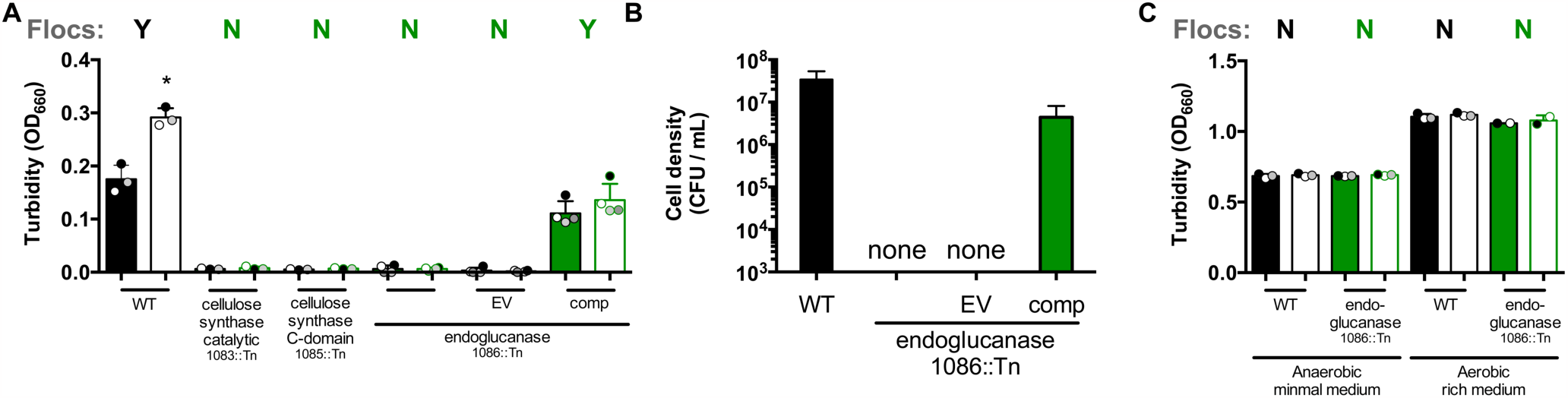
Cellulose synthase genes are required for survival in an aerobic minimal medium. (A, C) Turbidity of cultures after gentle vortexing (filled; 5 s, level 3) and after disruption of flocs by harsh vortexing (open; 10 pulses, level 10). Shaded circles indicate the same culture pre- and post-disruption. Y/N indicates visual observation of flocs (Y) or not (N). *Significant difference from the corresponding filled bar (P-value < 0.1) from a one-way ANOVA with Sidak’s multiple comparison posttest. WT, wild-type ZM4; EV, empty vector pSRKTc; comp, complementation vector pSRKTc-1086. (A) Cultures in aerobic minimal medium, ZYMM, inoculated from harshly vortexed cultures from conditions used in Fig. 3. (B) Colony forming units from cultures in panel A plated on PYG agar. ‘None’ indicates that no colonies were observed in 10 µl of a 10^-1^ dilution. (C) Cultures in anaerobic ZYMM or aerobic PYG medium inoculated from colonies. (A-C) Error bars, SD.

#### Respiratory NADH dehydrogenase is important for survival in an aerobic minimal medium

We questioned whether floc formation was stimulated by the aerobic environment or some aspect of aerobic respiration. NADH DH, encoded by ZMO1113, is the primary enzyme by which electrons enter the aerobic electron transfer chain in *Z. mobilis* (16–18). When cultured in an aerobic rich medium, *Z. mobilis* NADH DH mutants have been reported behave as if they are under anaerobic conditions, exhibiting improved growth and a high ethanol yield (16–18) instead of high levels of acetaldehyde and acetic acid typical of respiring *Z. mobilis* (19). We verified the improved growth trend and higher ethanol yield of three different NADH DH Tn mutants (1113::Tn) in aerobic PYG (Fig. 6). Expressing ZMO1113 from a plasmid in a 1113::Tn mutant resulted in a lower final OD and acetic acid and acetaldehyde accumulation in aerobic PYG similar to ZM4 (Fig. 6).

**Fig. 6.**
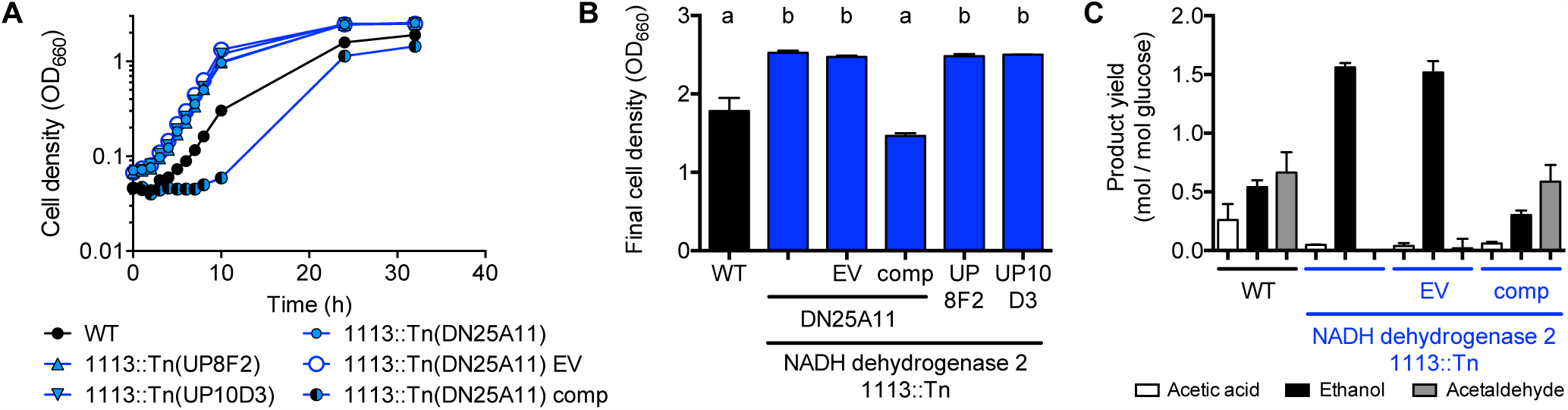
NADH DH mutants show improved growth trends and a higher ethanol yield in rich PYG medium. Representative growth curves (A), final OD values (B) and major fermentation product yields for wild-type ZM4 (WT) and NADH DH mutants (1113::Tn) in PYG medium. EV, empty vector pSRKTc; comp, complementation vector pSRKTc-1113. Error bars, SD; n=2. (A) Similar trends were observed for at least one other biological replicate.

Contrary to what we observed in PYG medium, the 1113::Tn mutants formed visible flocs in aerobic ZYMM, though to a lesser extent than in ZM4 cultures such that a change in OD was often negligible after disruption (Fig. 7A). Thus, floc formation is not solely a response to some aspect of NADH DH activity but is likely influenced by it. However, unlike in aerobic PYG, 1113::Tn culture growth was less than that of ZM4 in aerobic ZYMM. This trend of poor growth was exaggerated upon transfer of disrupted 1113::Tn mutant cultures to fresh medium (Fig. 7B, C). Expression of ZMO1113 from a vector improved survival of the 1113::Tn mutant whereas an empty vector did not (Fig. 7). Thus, we conclude that while NADH DH activity is dispensable and perhaps even detrimental in an aerobic rich medium like PYG, it is important for survival in an aerobic minimal medium.

**Fig 7.**
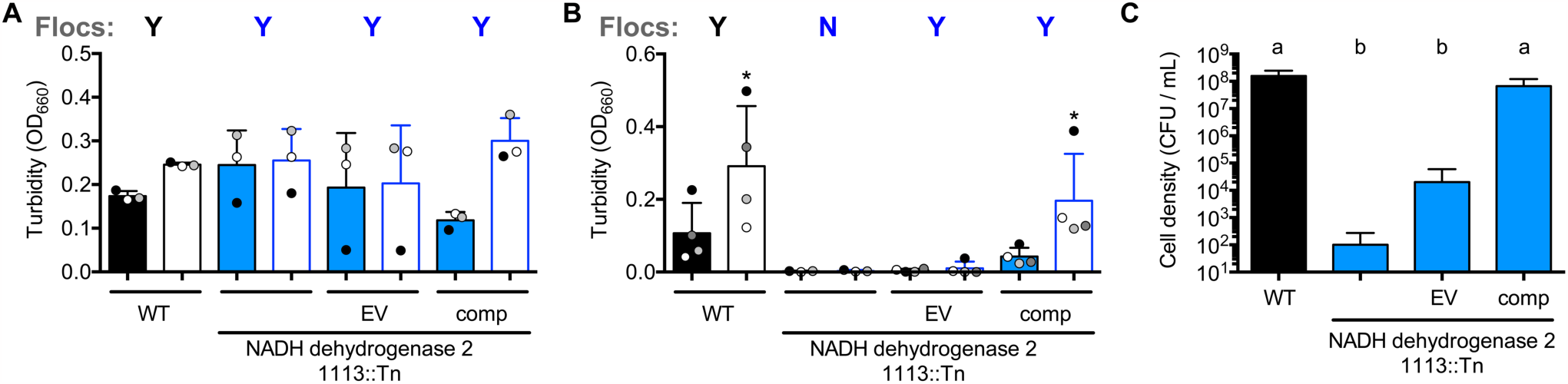
NADH DH is important for survival in an aerobic minimal medium. (A, B) Turbidity of cultures grown in an aerobic minimal medium (ZYMM) after gentle vortexing (filled; 5 s, level 3) and after disruption of flocs by harsh vortexing (open; 10 pulses, level 10). Cultures were inoculated from colonies (A) or inoculated from harshly vortexed cultures from conditions in panel A (B). Shaded circles indicate the same culture pre- and post-disruption. Y/N indicates visual observation of flocs (Y) or not (N). *Significant difference from the corresponding filled bar (P-value < 0.1) from a one-way ANOVA with Sidak’s multiple comparison posttest. (C) Colony forming units from cultures in panel B plated on PYG agar. Bars with different letters are significantly different (P-value < 0.) as determined by a one-way ANOVA with Sidak’s multiple comparison posttest. (A-C) Error bars, SD. EV: empty vector pSRKTc; comp: complementation vector pSRKTc-1113.

The poor viability of NADH DH mutants in aerobic ZYMM rules out several possibilities that could have explained the low viability of the flocculation mutants, 0919::Tn and 1086::Tn. First, since NADH DH mutants produce little acetic acid, flocculation mutant death was unlikely due to acidification from acetic acid accumulation. In further support of this notion, supplementing aerobic ZYMM with 100 mM MOPS, pH7, did not stimulate growth of 0919::Tn or 1086::Tn mutants, despite partially alleviating the pH drop (Fig. 8); ZM4, 0919::Tn, and 1086::Tn culture pH values reached ~4 without MOPS whereas pH values reached ~5.5 for ZM4 and ~6 for 0919::Tn and 1086::Tn with MOPS. Second, because NADH DH mutants produce little acetaldehyde, flocculation mutant death was unlikely due to acetaldehyde accumulation alone. We also tested whether acetaldehyde could be a factor that stimulates floc formation by adding acetaldehyde to cultures growing in anaerobic ZYMM, conditions in which acetaldehyde is normally not produced. The addition of acetaldehyde to anaerobic ZYMM resulted in a lower final OD but did not induce floc formation (Fig. 9).

**Fig. 8.**
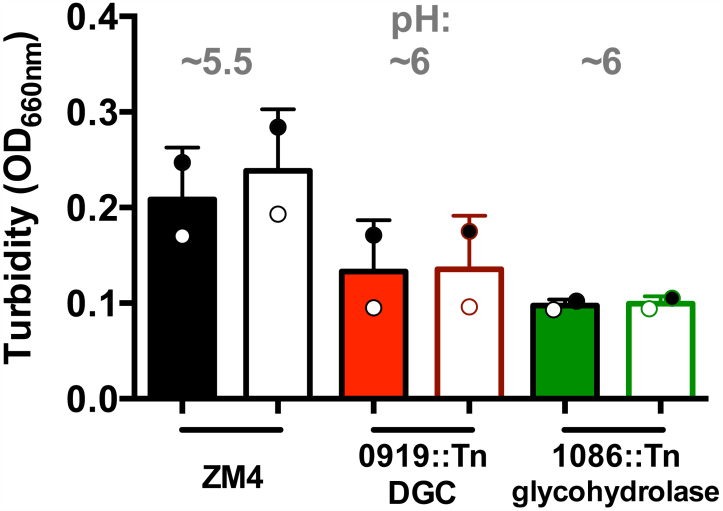
Adding MOPS buffer does not promote growth in an aerobic minimal medium. Turbidity of cultures grown in an aerobic minimal medium (ZYMM) after gentle vortexing (filled; 5 s, level 3) and after harsh vortexing (open; 10 pulses, level 10) to disrupt any flocs. Shaded circles indicate the same culture pre- and post-disruption.

**Fig. 9.**
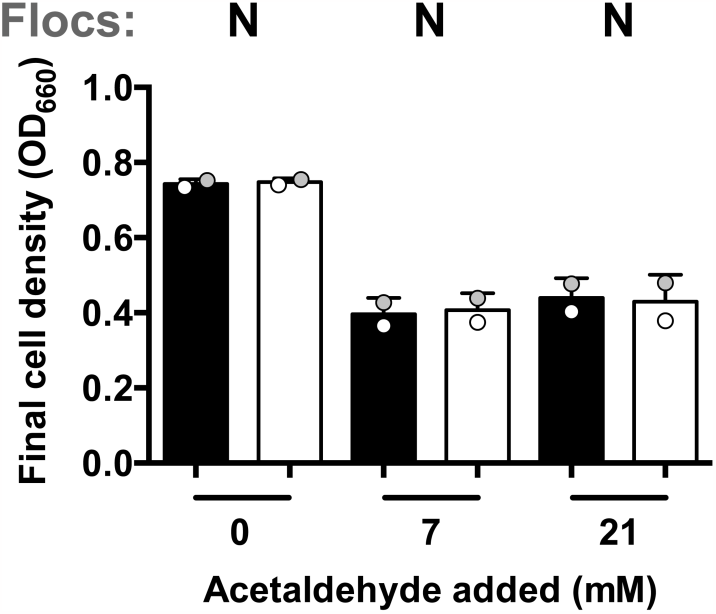
Acetaldehyde does not induce floc formation under anaerobic conditions. Turbidity of ZM4 cultures grown in an anaerobic minimal medium (ZYMM) with the indicated about of acetaldehyde after gentle vortexing (filled; 5 s, level 3) and after harsh vortexing (open; 10 pulses, level 10) to disrupt any flocs. Shaded circles indicate the same culture pre- and post-disruption. Y/N indicates visual observation of flocs (Y) or not (N).

#### Vitamins as a possible protective factors in yeast extract

Whereas ZM4 always formed flocs in aerobic ZYMM, we did not observe flocs when ZM4 was grown aerobically in PYG medium (Fig. 1). Many other studies on aerobically grown *Z. mobilis* used a medium with 0.5% yeast extract as the only undefined supplement (16, 17, 23-27). We therefore tested whether the addition of 0.5% yeast extract to ZYMM would improve growth and/or prevent floc formation. The addition of yeast extract averted the toxic effects of aerobic growth for all strains tested (Fig. 10A).H owever, the effect on floc formation in ZM4 cultures was variable; flocs were observed in ZM4 cultures inoculated from colonies, although cultures were mostly turbid, as seen from the negligible change in OD after harsh vortexing (Fig. 10A). In subsequent cultures inoculated from these harshly vortexed cultures, no flocs were visible to the eye (Fig. 10B). Similar effects on floc formation were observed for the 1113::Tn mutant (Fig10A, B).

**Fig. 10.**
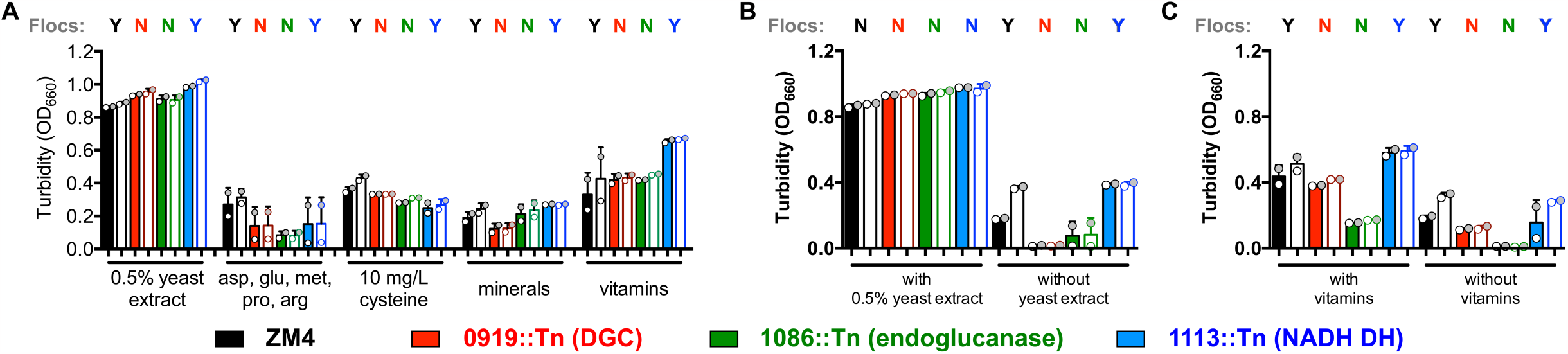
Yeast extract and vitamins promote growth of oxygen-sensitive mutants. Turbidity of cultures grown in an aerobic minimal medium (ZYMM) with the indicated supplement after gentle vortexing (filled; 5 s, level 3) and after disruption of flocs by harsh vortexing (open; 10 pulses, level 10). Cultures were inoculated from colonies (A) or inoculated from harshly vortexed cultures from panel A (B, C). Shaded circles indicate the same culture pre- and post-disruption. Y/N indicates visual observation of flocs (Y) or not (N).

We then screened several possibilities for why supplements like yeast extract might stimulate aerobic growth. Specifically we tested amino acids with antioxidant properties (28–32), mineral availability, and vitamins. The amino acid supplement was based on the expected concentration of arginine, aspartate, glutamate, methionine, and proline in yeast extract (33). The mixed amino acid supplement did not improve growth trends, nor did increasing the mineral concentration 10-fold over what is normally provided in ZYMM (Fig. 10A). Cysteine is a potent antioxidant and is commonly used to scavenge O_2_ in anaerobic media (32). Although not listed as a component of yeast extract, we tested a cysteine supplement based on the expected concentration of oxidized cystine in yeast extract (33). The cysteine supplement appeared to have a mild stimulatory effect on aerobic growth in ZYMM compared to adding the amino acid mixture or a higher concentration of minerals (Fig. 10A). Yeast extract is also rich in B vitamins, some of which can act as redox mediators (34). We therefore tested the addition of a readily available vitamin mixture (35) with B vitamins within the range of what is typical of yeast extract (34). The vitamin supplement stimulated aerobic growth of all strains, notably with the 1113::Tn mutant showing a higher final OD, similar to what was observed in aerobic PYG (Fig. 10a vs. Fig. 6B).

To determine if vitamins improved survival of oxygen-sensitive mutants, we then transferred a 1% inoculum of stationary phase-cultures to fresh identical medium with and without the supplements. While all strains grew to a higher final OD with vitamins than without, the two flocculation mutants did not grow as much as when inoculated from colonies (Fig. 10C vs Fig. 10A). Thus, while the vitamin supplement promoted aerobic growth of all strains, it did not appear to completely protect against the toxic aspects of aerobic growth.

We attempted to gain further insight into what conditions might protect *Z. mobilis* during aerobic growth by examining a chemogenomic profiling database in which the fitness effect from Tn disruptions of nearly every *Z. mobilis* ZM4 gene was tested across hundreds of conditions, including aerobic and anaerobic conditions with and without undefined supplements (36). However, none of the mutants examined in our study showed a significant growth defect during aerobic growth in a defined medium in the chemogenomic profiling database (36) (Fig. 11). In fact, the cellulose synthase gene cluster, ZMO1083-6 Tn mutants were reported to generally have positive fitness values across all conditions (36). The database also did not agree with predictions from our study and others (16–18) that an interruption in ZMO1113 encoding NADH DH would result in a fitness advantage in rich aerobic media (36). We speculate that these discrepancies are due to differences in the way the experiments were performed. For example, the chemogenomic profiling was carried out in 10-ml volumes (36), which might have limited aeration compared to the 5 ml volumes we used. Also, the defined medium used in the chemogenomic profiling contained four different B vitamins (36), which we showed can limit the toxic effects of aerobic growth (Fig. 10). Another possibility is that growing the entire library of mutants in the same test tube for chemogenomic profiling could have led to cross-complementation, rescuing mutants that would otherwise be subject to the toxic aspects of aerobic growth if grown as a clonal population.

**Fig. 11.**
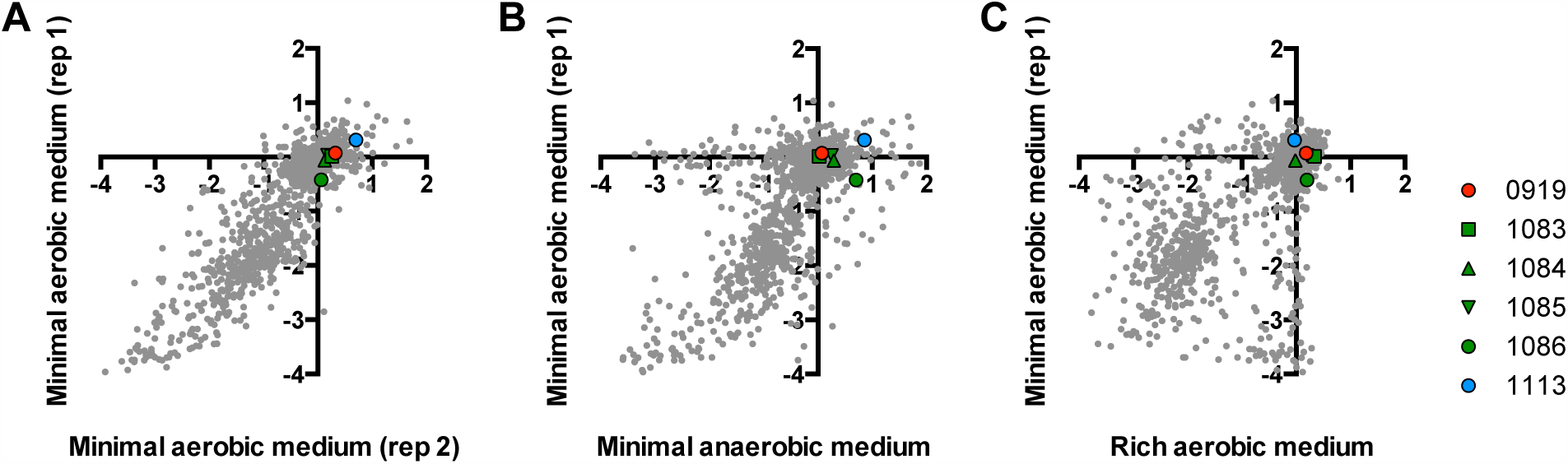
Comparison of gene fitness values from a chemogenomic profiling study (36). X and Y-axis values are gene fitness values for the indicated growth condition. Gene fitness values reflect the change in population frequency for a given Tn mutant from the time of inoculation until the end of the experiment (6-8 generations). Negative values indicate that a Tn insertion lowered strain fitness. Positive values indicate that a Tn insertion increased strain fitness. More information on the conditions and calculations used to determine gene fitness values is available elsewhere (36, 39). Each grey dot is the gene fitness value for a given ZM4 gene. Each graph compares gene fitness values in an aerobic minimal medium (rep 1) with a replicate in an identical growth condition (A; control; rep 2), an anaerobic minimal medium (B) or an aerobic rich medium (C).

#### Discussion

We have demonstrated that the ability to form flocs and the aerobic respiration enzyme NADH DH are important for *Z. mobilis* ZM4 survival in an aerobic defined medium. Floc formation and respiration could work in concert; by concentrating respiratory activities, the community of cells in a floc might create a protective anaerobic microenvironment. The importance of cells clustering might also help explain why flocculation mutants grew from colonies but not when transferred to fresh medium (Fig. 5 and 10). However, in opposition to this notion of synergistic activities, it is not obvious why starting respiration-deficient NADH DH mutant cultures from colonies would promote growth (Fig. 7). Rather, the formation of flocs and initial growth by an NADH DH mutant suggests that flocs might have protective properties independent of respiration. Such protection could be similar to that afforded by exopolysaccharide secreted by *Leuconostoc mesenteroides,* which was shown to protect cells from reactive oxygen by lowering the dissolved O_2_ concentration, although the biochemical mechanism by which this occurs remains unknown (37). One possible contributing factor to the poor growth of the NADH DH mutant upon transfer could stem from having transferred cultures after turbidity had leveled off, indicating a cessation of growth. There is evidence that NADH DH activity is important for survival in starved cultures, periodically fed glucose (23). Thus, a disadvantage under starvation conditions could contribute to an explanation for why NADH DH mutants grew poorly upon transfer.

Undefined supplements promoted growth of all strains tested and in some cases inhibited floc formation (Fig. 1 and 10). We determined that mineral availability, anti-oxidant amino acids, and vitamins by themselves cannot fully explain why supplements like yeast extract have a protective role (Fig. 10). One possibility is that yeast extract contributes to a reduced environment as yeast extract has been shown to lower the oxidation-reduction potential of media (34); components in yeast extract could help to keep critical cell components in a reduced state during aerobic growth. Redox-active B vitamins in yeast extract, could contribute to this role (34, 38), and could also help explain why adding vitamins alone promoted aerobic growth in ZYMM (Fig. 10). Another possibility is that yeast extract and vitamins provides a critical nutrient that *Z. mobilis* has difficulty synthesizing in an aerobic environment, perhaps due to an O_2_-sensitive enzyme.

Use of yeast extract in studies of *Z. mobilis* aerobic respiration have also led to observations by others that NADH DH activity, and respiration in general, is dispensable and even comes with a fitness cost during growth (16-18, 25). These observations raise questions as to how genes that seemingly come with a fitness cost, like that encoding NADH DH, have been maintained in *Z. mobilis*. Our observations showing that NADH DH is important for survival in an aerobic defined medium (Fig. 7) suggest that aerobic respiration is important for survival in natural environments that are deficient in protective factors like those found in yeast extract.

The exact mechanism of toxicity and signal for flocculation in aerobic conditions remains elusive. Nonetheless, we have uncovered general conditions and key genetic factors responsible flocculation and survival. This knowledge could be used to control flocculation and potentially promote survival in response to stress conditions in industrial hydrolysates.

### Methods

#### Strains and growth conditions

All strains are described in Table 1. *Zymomonas mobilis* ZM4 (ATCC 31821) and all Tn mutants were provided by J. M. Skerker and A. P. Arkin, UC Berkeley (39). When only one Tn mutant was used, we selected the mutant with the insertion site closest to the N-terminus. Exact insertion site locations are listed elsewhere (39). Transposon insertions were verified by PCR and sequencing of PCR products. Strains were streaked from frozen 25% glycerol stocks preserved at −80°C onto PYG agar (2% peptone, 1% yeast extract, 2% glucose, 1.5% agar) with antibiotics as appropriate. For growth experiments, colonies were then inoculated to either 5 ml aerobic or 10 ml anaerobic liquid media and incubated at 30°C with shaking at 225 rpm. ZYMM with 50mM glucose, 100 nM calcium pantothenate and 10 mM NH_4_Cl was used as the minimal medium (4). Where indicated, cultures were also supplemented with a mixture of amino acids with final concentrations of 0.1 mg/ml for aspartate, arginine, methionine, and proline, and 0.6 mg/ml potassium glutamate. The cysteine-HCl supplement was added to a final concentration of 0.01 mg/ml. The mineral supplement contained 10-fold more of the mineral supplement typically added to ZYMM (4). The vitamin supplement was added from a 100X stock solution described elsewhere (35). Unless stated otherwise, aerobic test tubes had loose-fitting caps to allow for air exchange. For anaerobic conditions, media was bubbled with N_2_ gas and then tubes were sealed with rubber stoppers (Geo-Microbial Technologies, Ochelata, OK) and aluminum crimps. *Escherichia coli* strains used for cloning were grown in LB broth or on LB agar. Where appropriate, tetracycline was used at 5 μg/ml for *Z. mobilis* and at 15 μg/ml for *E. coli* and kanamycin was used at 100 μg/ml for *Z. mobilis*.

**Table 1.**
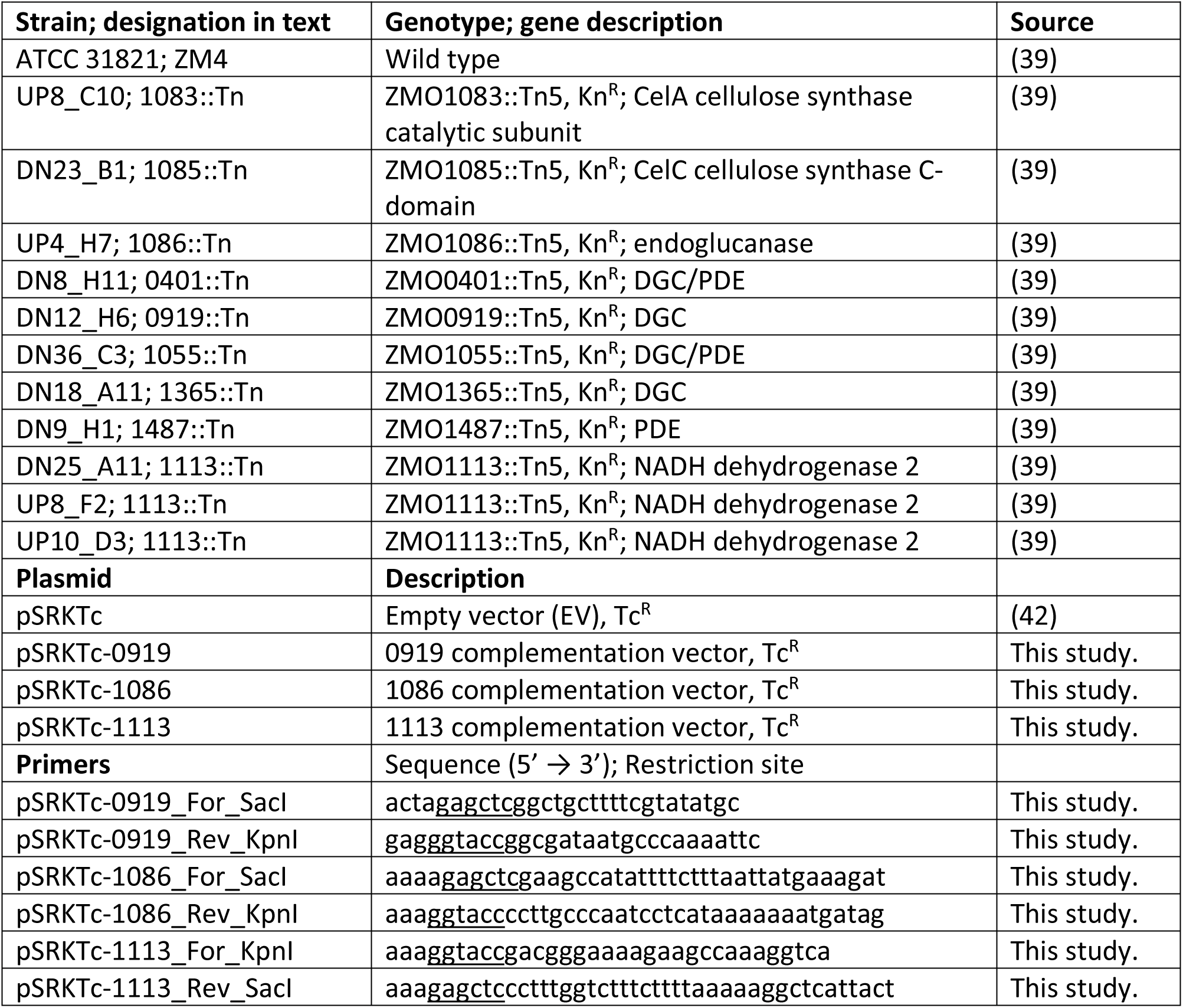
**Strains, plasmids and primers.** Kn^R^, kanamycin resistance cassette; Tc^R^, tetracycline resistance cassette; DGC, diguanylate cyclase; PDE, phosphodiesterase; Tn5, transposon Tn5.

#### Construction of complementation vectors

All plasmids and primers are described in Table 1. All enzymes and competent cells were used according to the manufacturer’s instructions. Primers were designed to amplify entire genes and promoter regions as predicted using the algorithm BPROM (www.softberry.com) (40) and to introduce flanking ScaI and KpnI restriction sites. PCR products were then digested and ligated into pSRKTc that had been cut with the same enzymes. The ligation reaction was transformed into *E. coli* NEB10β (NEB) and plated on LB agar with tetracycline. Transformants were screened by PCR and verified by sequencing. Purified plasmids were then transformed into electrocompetent cells of the appropriate *Z. mobilis* strain as described (41).

#### Cellulase and mechanical disruption of flocs

All vortexing levels refer to that for a VWR Mini Vortexer. Pre-disruption turbidity was assessed after vortexing cultures at level 3 for 5 seconds. To enzymatically disrupt flocs, 50 μL of *Aspergillus niger* cellulase (MP; 1 % w/v) in 0.1 M MES Buffer, pH 3.5 was added per 1 mL of culture and incubated overnight at 30°C. Treated cultures were then vortexed at for 5 seconds at speed 6 prior to measuring OD_660_ measurement. To mechanically disrupt flocs, cultures were vortexed with 10 pulses at level 10. Disruption was performed after the turbid fraction had reached a maximum OD_660_ value, typically within six days, although sometimes longer for slow-growing strains harboring plasmids. Culture turbidity was assayed by optical density at 660 nm (OD_660_) using a Genesys 20 visible spectrophotometer (Thermo-Fisher, Pittsburgh, PA).

#### Enzymatic characterization of the extracellular matrix

Enzymes were added to flocs grown in aerobic ZYMM then added to each test tube and incubated at 30°C with shaking at 225 rpm. Cultures were periodically resuspended by vortexing at level 3 for 5 seconds and optical density readings taken. Stock solutions of enzymes were prepared in water except for cellulase which was prepared in 0.1 M MES, pH 3.5. Final concentrations were: DNase, 20 μg/ml; proteinase K, 800 μg/ml; cellulase 100 μg/ml.

#### Scanning electron microscopy

Flocculent culture was added to a 0.1% poly-L-lysine treated glass coverslip and incubated at room temperature for 5 min. Cells were then fixed with 3% glutaraldehyde in phosphate buffered saline (PBS) and incubated at 4°C for 1 h. Fixative was removed with three washes of 4°C PBS. The sample was dehydrated using a graded ethanol series (30, 50, 70, 90,and 95%), at 4°C for 5 min at each concentration, followed by three changes of 100% ethanol at room temperature for 5 min each. The sample was thin critical point dried using CO_2_ in a Balzers CPD 030 critical point drier. The dried sample was then placed on an aluminum stub, sputter coated with gold/palladium (60:40), and imaged using a Jeol JSM-5800 scanning electron microscope at the Indiana University Electron Microscopy Center.

#### Statistics

All statistical analyses were performed using Prism 6.0h (Graphpad Software, Inc.)

## Acknowledgements

This work was funded by the Indiana University College of Arts and Sciences. JM was supported in part during by a National Science Foundation CAREER award # MCB-1749489.

We are grateful to JM Skerker and AP Arkin for providing the transposon mutants and parental strain. We thank BD Stein at the Indiana University Electron Microscopy Center and AL McCully for the electron microscopy image taken during a 2015 Z620 graduate course. We thank Jeffrey Mazny with assistance with experiments that helped shape the study.

